# Antibody binding to SARS-CoV-2 S glycoprotein correlates with, but does not predict neutralization

**DOI:** 10.1101/2020.09.08.287482

**Authors:** Shilei Ding, Annemarie Laumaea, Romain Gasser, Halima Medjahed, Marie Pancera, Leonidas Stamatatos, Andrew McGuire, Renée Bazin, Andrés Finzi

## Abstract

Convalescent plasma from SARS-CoV-2 infected individuals and monoclonal antibodies were shown to potently neutralize viral and pseudoviral particles carrying the S glycoprotein. However, a non-negligent proportion of plasma samples from infected individuals as well as S-specific monoclonal antibodies were reported to be non-neutralizing despite efficient interaction with the S glycoprotein in different biochemical assays using soluble recombinant forms of S or when expressed at the cell surface. How neutralization relates to binding of S glycoprotein in the context of viral particles remains to be established. Here we developed a pseudovirus capture assay (VCA) to measure the capacity of plasma samples or antibodies immobilized on ELISA plates to bind to membrane-bound S glycoproteins from SARS-CoV-2 expressed at the surface of lentiviral particles. By performing VCA and neutralization assays we observed a strong correlation between these two parameters. However, while we found that plasma samples unable to capture viral particles did not neutralize, capture did not guarantee neutralization, indicating that the capacity of antibodies to bind to the S glycoprotein at the surface of viral particles is required but not sufficient to mediate neutralization. Altogether, our results highlights the importance of better understanding the inactivation of S by plasma and neutralizing antibodies.

## Introduction

The etiologic agent of the Coronavirus Disease (COVID-19), the severe acute respiratory syndrome coronavirus 2 (SARS-CoV-2), spread from China to the rest of the world and is the cause of the current pandemic [1]. In the absence of an effective vaccine to prevent SARS-CoV-2 infection, alternative approaches to treat or prevent acute COVID-19 are desperately needed. A promising approach is the use of convalescent plasma containing anti-SARS-CoV-2 antibodies collected from donors who have recovered from COVID-19 [2]. Convalescent plasma therapy was successfully used in the treatment of SARS, MERS and influenza H1N1 pandemics and was associated with improvement of clinical outcomes [3-5]. The transfer of convalescent plasma to COVID-19 patients was shown to be well tolerated and presented some positive signs [6-10]. Similarly, some antibodies targeting the Spike (S) glycoprotein from SARS-CoV-2, isolated from virus infected individuals, were shown to potently neutralize viral and pseudoviral particles carrying the S glycoprotein of the virus [11-16] and in some instances protected small animals from SARS-CoV-2 infection [17, 18].

Nonetheless, a non-negligent proportion of plasma samples from infected individuals as well as S-specific monoclonal antibodies were reported to be non-neutralizing despite efficient interaction with the S glycoprotein in different biochemical assays using soluble recombinant forms of S or when expressed at the cell surface [12, 16]. Thus, raising the question of how neutralization relates to binding of S in the context of viral or pseudoviral particles. To address this question, we adapted a previously-described virus capture assay (VCA)[19] to measure the capacity of plasma samples and monoclonal antibodies to bind to membrane-bound S glycoproteins from SARS-CoV-2 expressed at the surface of lentiviral particles and compared to their neutralization activity.

## Materials and Methods

### Ethics statement

All subjects gave their informed consent for inclusion before they participated in the study. The study was conducted in accordance with the Declaration of Helsinki, and the protocol was approved by the Ethics Committee of CHUM (19.381, approved on March 25, 2020). Donors met all donor eligibility criteria: previous confirmed COVID-19 infection and complete resolution of symptoms for at least 14 days.

### Plasmids

The plasmids expressing the human coronavirus Spike of SARS-CoV-2 and SARS-CoV-1 were kindly provided by Stepfan Pöhlman and were previously reported [20]. The pNL4.3 R-E-Luc was obtained from NIH AIDS Reagent Program. The vesicular stomatitis virus G (VSV-G)-encoding plasmid (pSVCMV-IN-VSV-G) was previously described [14].

### Cell lines

293T human embryonic kidney cells (obtained from ATCC) and Cf2Th cells (kind gift from Joseph Sodroski, DFCI) were maintained at 37°C under 5% CO2 in Dulbecco’s modified Eagle’s medium (DMEM) (Wisent) containing 5% fetal bovine serum (VWR), 100 UI/ml of penicillin and 100μg/ml of streptomycin (Wisent). The 293T-ACE2 cell line was previously reported [14].

### Virus capture assay

The assay was modified from a previous published method [19]. Briefly, pseudoviral particles were produced by transfecting 2×10^6^ HEK293T cells with pNL4.3 Luc R-E-(3.5μg), plasmids encoding for SARS-CoV-2 or SARS-CoV-2 Spike (3.5μg) glycoproteins using the standard calcium phosphate protocol. Forty-eight hours later, supernatant-containing virion was collected and cell debris was removed through centrifugation (1,500 rpm for 10 min). To immobilize plasma on ELISA plates, white MaxiSorp ELISA plates (Thermo Fisher Scientific) were incubated with 1:500 diluted plasma in 100μl phosphate-buffered saline (PBS) overnight at 4°C. Unbound antibodies or plasma were removed by washing the plates twice with PBS. Plates were subsequently blocked with 3% bovine serum albumin (BSA) in PBS for 1 h at room temperature. After two washes with PBS, 200μl of virus-containing supernatant was added to the wells. Viral capture by any given plasma sample was visualized by adding 10×10^4^ SARS-CoV-2-resistant Cf2Th cells in full DMEM medium per well. Forty-eight hours post-infection, cells were lysed by the addition of 30μl of passive lysis buffer (Promega) and three freeze-thaw cycles. An LB941 TriStar luminometer (Berthold Technologies) was used to measure the luciferase activity of each well after the addition of 100μl of luciferin buffer (15 mM MgSO_4_, 15 mM KH_2_PO_4_ [pH 7.8], 1 mM ATP, and 1mM dithiothreitol) and 50μl of 1mM D-luciferin potassium salt (Prolume).

### Virus neutralization assay

Target cells were infected with single-round luciferase-expressing lentiviral particles as described previously [14]. Briefly, 293T cells were transfected by the calcium phosphate method with the lentiviral vector pNL4.3 R-E-Luc (NIH AIDS Reagent Program) and a plasmid encoding for SARS-CoV-2 Spike, SARS-CoV-1 Spike at a ratio of 5:4. Two days post-transfection, cell supernatants were harvested and stored at –80°C until use. 293T-ACE2 target cells were seeded at a density of 1×10^4^ cells/well in 96-well luminometer-compatible tissue culture plates (Perkin Elmer) 24h before infection. Recombinant viruses in a final volume of 100μl were incubated with the indicated sera dilutions (1/50; 1/250; 1/1250; 1/6250; 1/31250) for 1h at 37°C and were then added to the target cells followed by incubation for 48h at 37°C; cells were lysed by the addition of 30μl of passive lysis buffer (Promega) followed by one freeze-thaw cycle. An LB941 TriStar luminometer (Berthold Technologies) was used to measure the luciferase activity of each well after the addition of 100μl of luciferin buffer (15mM MgSO_4_, 15mM KPO_4_ [pH 7.8], 1mM ATP, and 1mM dithiothreitol) and 50μl of 1mM d-luciferin potassium salt (Prolume). The neutralization half-maximal inhibitory dilution (ID_50_) or the neutralization 80% inhibitory dilution (ID_80_) represents the sera dilution to inhibit 50% or 80% of the infection of 293T-ACE2 cells by recombinant viruses bearing the indicated surface glycoproteins.

## Results and Discussion

### Virus capture assay

To measure the capacity of plasma samples and monoclonal antibodies immobilized on ELISA plates to bind to membrane-bound S glycoproteins from SARS-CoV-2 expressed at the surface of lentiviral particles we adapted a previously-described virus capture assay (VCA)[19]. The pseudoviral particles used in this assay are generated by transfecting HEK293T cell with the pNL4.3 Nef-Luc Env-construct [21-24]. This construct is co-transfected with a plasmid encoding the S glycoprotein from SARS-CoV-2 and a plasmid encoding the G glycoprotein from vesicular stomatitis virus (VSV-G), resulting in a virus capable of a single round of infection. Pseudovirus-containing supernatants are added to plasma or antibody-coated ELISA plates and unbound pseudoviruses are washed away. Retention of pseudoviruses on the surface of the plate by S-specific antibodies is visualized by the addition of Cf2Th cells. These cells are refractory to SARS-CoV-2 S-mediated entry (not shown) but are infected by the bound pseudoparticles via the incorporated G glycoprotein from VSV. Cf2Th infection is measured as a function of luciferase activity two days later. A scheme of the assay is depicted in Figure 1A. This assay was used with plasma samples from SARS-CoV-2-convalescent individuals and, as expected, showed a robust capture of pseudoviral particles whereas plasma obtained from SARS-CoV-2 uninfected individuals failed to do so (Figure 1B). Similarly, previously-described SARS-CoV-2 S-specific monoclonal antibodies isolated from convalescent donors [12] were able to capture pseudoviruses expressing the S glycoprotein from SARS-CoV-2. Of note, the capture was specific since plasma from convalescent individuals and monoclonal antibodies failed to capture similar pseudoviruses expressing the S glycoprotein from SARS-CoV-1 (Figure 1B).

**Figure 1.**
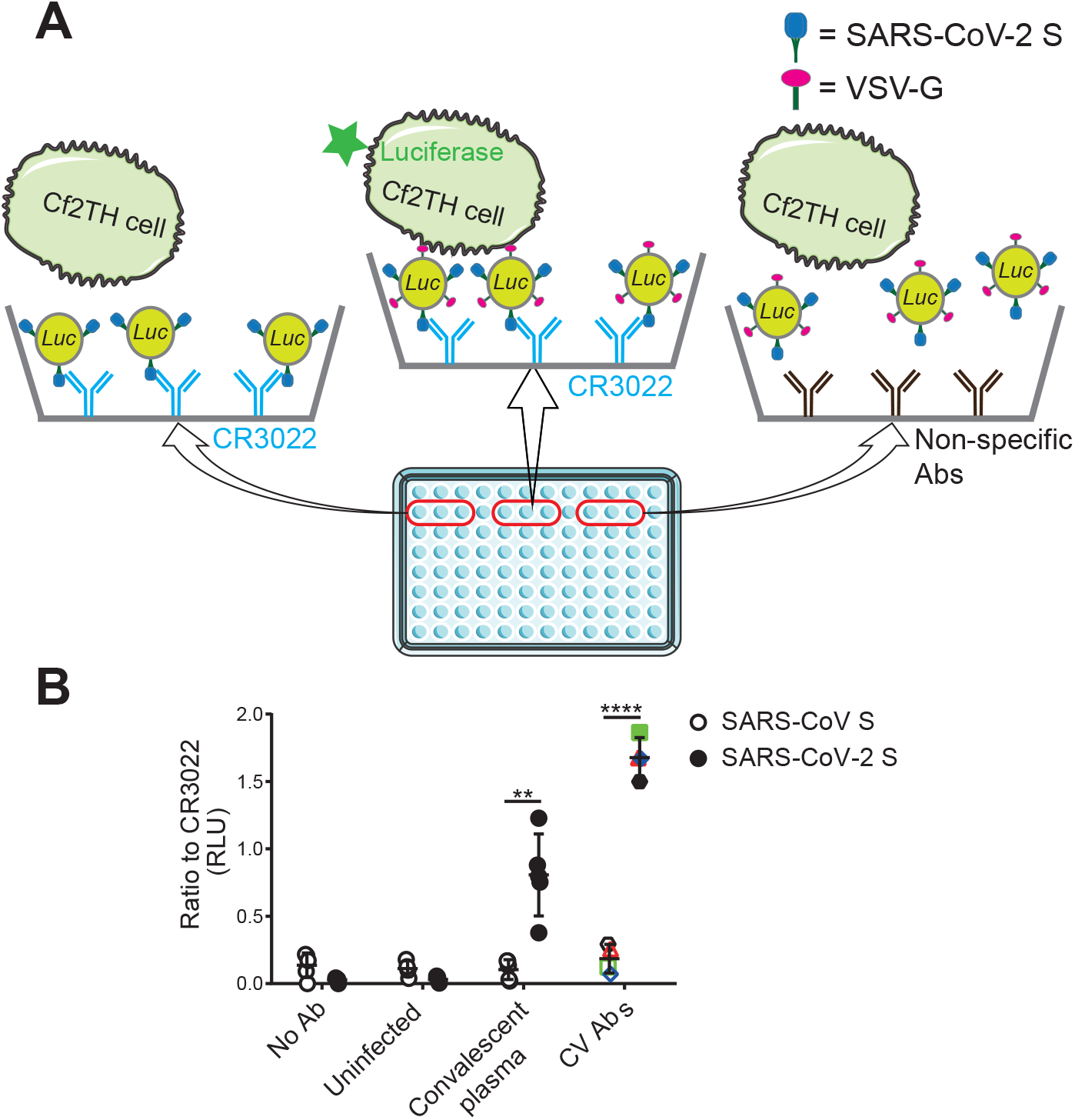
Depiction of the Virus capture assay. (**A**) As shown in the scheme, 96-well ELISA plates were coated with SARS-CoV-2 S specific monoclonal antibodies or plasma recovered from SARS-CoV-2 uninfected individuals (uninfected) or plasma recovered six weeks after symptoms onset (convalescent plasma). Viral particles encoding for luciferase and bearing VSV-G glycoprotein and SARS-CoV-1 S or SARS-CoV-2 S glycoproteins were added to the wells. Free virions were washed away and Cf2Th cells which are refractory to SARS-CoV-2 S-mediated entry were added to the wells. After 48 hours, cells were lysed and luciferase activity was measured. (**B**) Previously-described SARS-CoV-2 S specific monoclonal antibodies [12] (CV1 -black hexagon, CV2 – red triangle, CV24 – green square and CV30 – blue diamond), plasma recovered from SARS-CoV-2 uninfected individuals (uninfected) or plasma recovered six weeks after symptoms onset (convalescent plasma) were tested for the binding with viral particles bearing SARS-CoV-1 S (hollow) or SARS-CoV-2 S glycoproteins (solid). Relative light unit (RLU) obtained from CR3022 was used as control (set as one). Data shown are the mean ± standard deviation (SD) of three independent experiments performed in triplicate. Statistical significance was evaluated using a paired t test (**, P<0.01, ****, P<0.0001).

### Recognition of S glycoproteins at the surface of pseudoviral particles is required but no sufficient to neutralize

It is presently unclear whether the capacity of plasma from convalescent donors or monoclonal antibodies to neutralize pseudoviral particles correlates with their capture efficiency. Therefore, we first measured the capacity of plasma samples recovered six weeks after symptoms onset from twenty-five convalescent individuals to neutralize pseudoparticles bearing the SARS-CoV-2 S glycoprotein using 293T cells stably expressing ACE2 as target cells, as described [14]. Neutralizing activity, as defined by the neutralization half-maximum inhibitory dilution (ID_50_), is shown in Figure 2A. These results illustrate the variable capacity of different convalescent plasma samples to neutralize. We then measured the capacity of the same plasma samples to capture viral particles. As mentioned above, HEK293T cells were co-transfected with pNL4.3 Nef-Luc Env-together with plasmids expressing the SARS-CoV-2 S glycoprotein and VSV-G; released viral particles were collected two days after transfection. For the VCA, 96-well microplates were coated with plasma recovered from SARS-CoV-2 uninfected individuals (control) or plasma from convalescent donors or with the receptor-binding domain (RBD)-specific CR3022 monoclonal antibody. This antibody was added to each plate for normalization purposes. Pseudoviral particles were added to the plates and incubated for 4 hours at 37°C; the plates were then washed to remove unbound pseudoviruses. Cf2th cells were added to the wells, incubated at 37°C and lysed 48 hours later to measure luciferase activity. As reported in Figure 2B, while a few plasma samples from convalescent donors failed to capture viral particles, similar to plasma from uninfected donors, most did. Interestingly, even though we observed a significant correlation between virus capture and neutralization (Figure 2C), virus capture did not always translate into neutralization. Indeed, we observed several convalescent plasma samples that were able to efficiently capture pseudoviral particles but did not neutralize, thus indicating that while interaction of SARS-CoV-2 S glycoprotein is required for virus neutralization, it is not sufficient *per se*.

**Figure 2.**
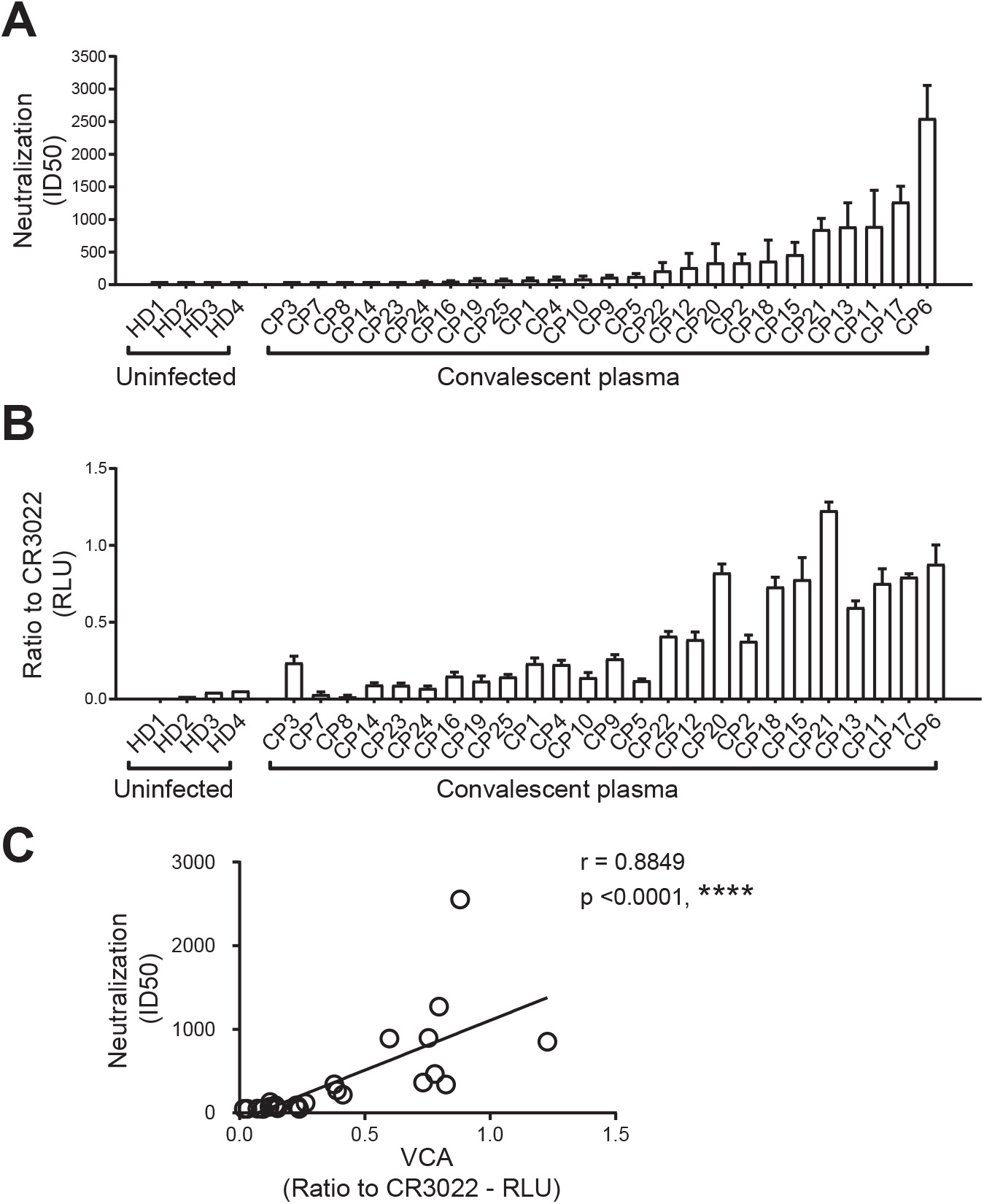
Recognition of S glycoproteins at the surface of pseudoviral particles is required but no sufficient to neutralize. (**A**) Pseudoviral particles coding for the luciferase reporter gene and bearing the SARS-CoV-2 S glycoprotein were used to infect 293T-ACE2 cells. Pseudoviruses were incubated with serial dilutions of plasma recovered from SARS-CoV-2 uninfected individuals (uninfected) or plasma recovered six weeks after symptoms onset (convalescent plasma) at 37°C for 1h prior to infection of 293T-ACE2 cells. (**A**) Neutralization half maximal inhibitory serum dilution (ID_50_) was determined using a normalized non-linear regression using Graphpad Prism software. (**B**) VSV-G-pseudotyped viral particles expressing the SARS-CoV-2 S glycoprotein were added to plates coated with plasma samples. Free virions were washed away and Cf2th cells were added to the wells. After 48 hours, cells were lysed and luciferase activity was measured. Luciferase signals were normalized to those obtained with the RBD-specific CR3022 antibody. Data shown are the mean ± SD of three independent experiments performed in triplicate. (**C**) Neutralization potency correlates with the ability of plasma samples to capture pseudoviral particles. Statistical significance was established with Spearman rank correlation test.

### The capacity of convalescent plasma to bind to the S glycoprotein of SARS-CoV-2 and neutralize pseudoviral particles decreases over time

Several reports described a significant decrease in the neutralization capacity of plasma from convalescent individuals starting six weeks after symptoms onset [13, 14, 25, 26]. To evaluate whether this was related to the capacity of plasma to capture viral particles, we analyzed the neutralization and viral capture capacity of serological samples obtained from fifteen convalescent donors at six and ten weeks after symptoms onset. As previously reported [13], we observed a significant decrease in the neutralizing capacity of samples over time (Figure 3A). A similar decrease in their capacity to capture viral particles was observed in the present study (Figure 3B). Accordingly, we noted a strong correlation between neutralization and virus capture (Figure 3C), suggesting that the decrease in neutralization over time might be due to the disappearance of antibodies able to recognize the S glycoprotein at the surface of viral particles.

**Figure 3.**
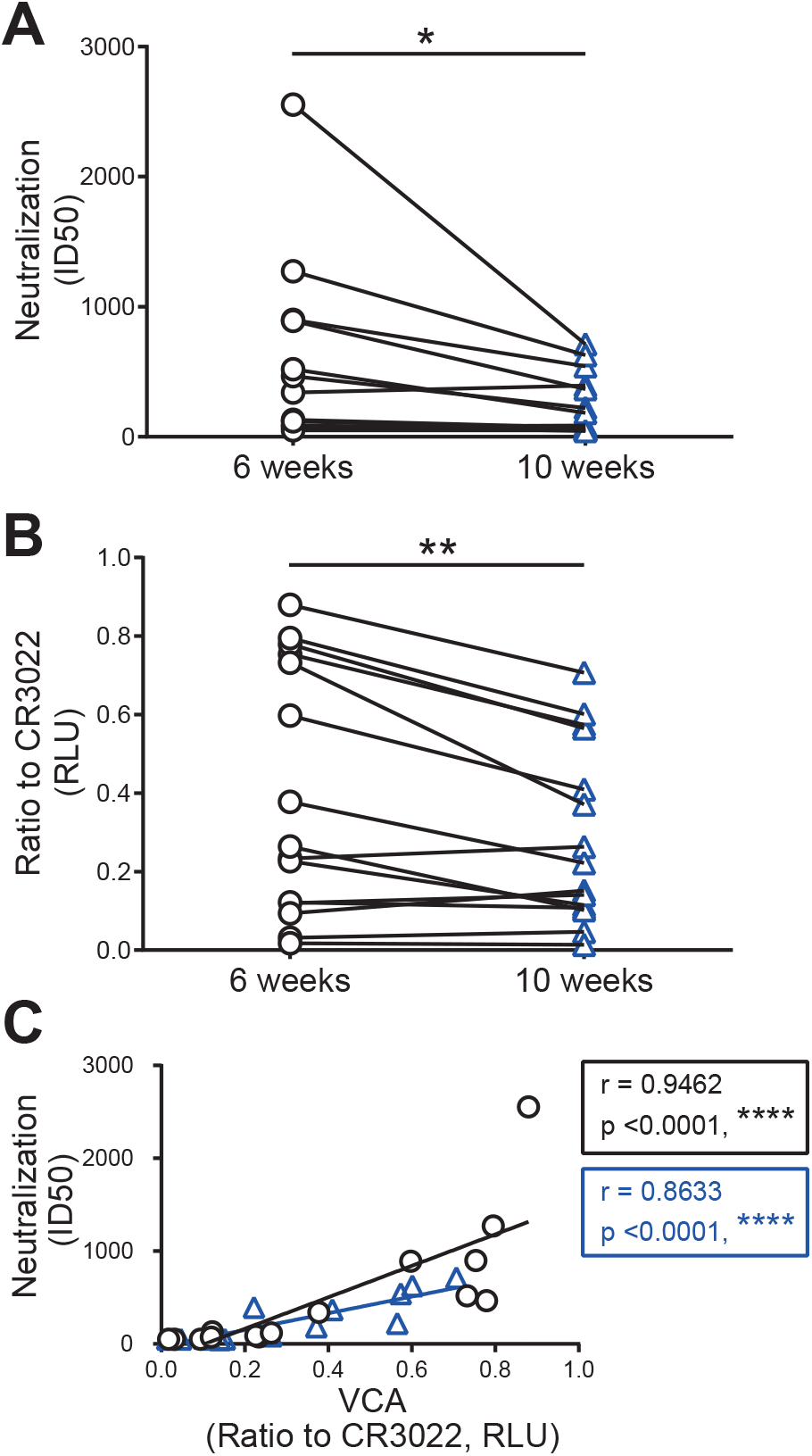
The capacity of convalescent plasma to bind to the S glycoprotein of SARS-CoV-2 and neutralize pseudoviral particles decreases over time. Pseudoviral particles coding for the luciferase reporter gene and bearing the SARS-CoV-2 S glycoprotein were used to infect 293T-ACE2 cells. Pseudoviruses were incubated with serial plasma dilutions at 37°C for 1h prior to infection of 293T-ACE2 cells. Convalescent plasma was recovered six (black circle) and ten (blue triangle) weeks after symptoms onset from fifteen individuals. (**A**) Neutralization half maximal inhibitory serum dilution (ID_50_) was determined using a normalized non-linear regression using Graphpad Prism software. (**B**) VSV-G-pseudotyped viral particles expressing the SARS-CoV-2 S glycoprotein were added to plates coated with plasma samples. Free virions were washed away and Cf2th cells were added to the wells. After 48 hours, cells were lysed and luciferase activity was measured. Luciferase signals were normalized to those obtained with the RBD-specific CR3022 antibody. Data shown are the mean ± SD of three independent experiments performed in triplicate. Statistical significance was evaluated using a paired t test (*, P<0.05, **, P<0.01). (**C**) Neutralization potency correlates with the ability of plasma samples to capture pseudoviral particles. Statistical significance was established with Spearman rank correlation test.

Altogether, using a newly designed virus capture assay, here we report that the capacity of antibodies to bind to the S glycoprotein at the surface of viral particles is required but not sufficient to mediate neutralization. Efforts to better understand the link between antibody interaction with S and virus neutralization might assist ongoing vaccine efforts aimed at eliciting neutralizing antibodies.

## Author Contributions

S.D. and A.F. conceived the study. S.D., A.L., R.G and HM. performed and interpreted the experiments. M.P., L.S. and A.M. contributed monoclonal antibodies. All the authors analyzed the data. S.D., R.B. and A.F. wrote the manuscript. Every author has read, edited and approved the final manuscript.

## Acknowledgments

The authors are grateful to the volunteers who donated plasma samples for this study and to Chantal Morrisseau and Laurie Gokool for sample collection. We also thank Dr M. Gordon Joyce (U.S. MHRP) for the monoclonal antibody CR3022.

## References

1. Zhou, P.; Yang, X. L.; Wang, X. G.; Hu, B.; Zhang, L.; Zhang, W.; Si, H. R.; Zhu, Y.; Li, B.; Huang, C. L.; Chen, H. D.; Chen, J.; Luo, Y.; Guo, H.; Jiang, R. D.; Liu, M. Q.; Chen, Y.; Shen, X. R.; Wang, X.; Zheng, X. S.; Zhao, K.; Chen, Q. J.; Deng, F.; Liu, L. L.; Yan, B.; Zhan, F. X.; Wang, Y. Y.; Xiao, G. F.; Shi, Z. L., A pneumonia outbreak associated with a new coronavirus of probable bat origin. Nature 2020, 579, (7798), 270–273.

2. Chen, L.; Xiong, J.; Bao, L.; Shi, Y., Convalescent plasma as a potential therapy for COVID-19. The Lancet infectious diseases 2020, 20, (4), 398–400.

3. Ko, J. H.; Seok, H.; Cho, S. Y.; Ha, Y. E.; Baek, J. Y.; Kim, S. H.; Kim, Y. J.; Park, J. K.; Chung, C. R.; Kang, E. S.; Cho, D.; Muller, M. A.; Drosten, C.; Kang, C. I.; Chung, D. R.; Song, J. H.; Peck, K. R., Challenges of convalescent plasma infusion therapy in Middle East respiratory coronavirus infection: a single centre experience. Antiviral therapy 2018, 23, (7), 617–622.

4. Hung, I. F.; To, K. K.; Lee, C. K.; Lee, K. L.; Chan, K.; Yan, W. W.; Liu, R.; Watt, C. L.; Chan, W. M.; Lai, K. Y.; Koo, C. K.; Buckley, T.; Chow, F. L.; Wong, K. K.; Chan, H. S.; Ching, C. K.; Tang, B. S.; Lau, C. C.; Li, I. W.; Liu, S. H.; Chan, K. H.; Lin, C. K.; Yuen, K. Y., Convalescent plasma treatment reduced mortality in patients with severe pandemic influenza A (H1N1) 2009 virus infection. Clinical infectious diseases : an official publication of the Infectious Diseases Society of America 2011, 52, (4), 447–56.

5. Cheng, Y.; Wong, R.; Soo, Y. O.; Wong, W. S.; Lee, C. K.; Ng, M. H.; Chan, P.; Wong, K. C.; Leung, C. B.; Cheng, G., Use of convalescent plasma therapy in SARS patients in Hong Kong. Eur J Clin Microbiol Infect Dis 2005, 24, (1), 44–6.

6. Bloch, E. M.; Shoham, S.; Casadevall, A.; Sachais, B. S.; Shaz, B.; Winters, J. L.; van Buskirk, C.; Grossman, B. J.; Joyner, M.; Henderson, J. P.; Pekosz, A.; Lau, B.; Wesolowski, A.; Katz, L.; Shan, H.; Auwaerter, P. G.; Thomas, D.; Sullivan, D. J.; Paneth, N.; Gehrie, E.; Spitalnik, S.; Hod, E. A.; Pollack, L.; Nicholson, W. T.; Pirofski, L. A.; Bailey, J. A.; Tobian, A. A., Deployment of convalescent plasma for the prevention and treatment of COVID-19. The Journal of clinical investigation 2020, 130, (6), 2757–2765.

7. Casadevall, A.; Joyner, M. J.; Pirofski, L. A., A Randomized Trial of Convalescent Plasma for COVID-19-Potentially Hopeful Signals. JAMA : the journal of the American Medical Association 2020.

8. Casadevall, A.; Pirofski, L. A., The convalescent sera option for containing COVID-19. The Journal of clinical investigation 2020, 130, (4), 1545–1548.

9. Joyner, M. J.; Wright, R. S.; Fairweather, D.; Senefeld, J. W.; Bruno, K. A.; Klassen, S. A.; Carter, R. E.; Klompas, A. M.; Wiggins, C. C.; Shepherd, J. R.; Rea, R. F.; Whelan, E. R.; Clayburn, A. J.; Spiegel, M. R.; Johnson, P. W.; Lesser, E. R.; Baker, S. E.; Larson, K. F.; Ripoll, J. G.; Andersen, K. J.; Hodge, D. O.; Kunze, K. L.; Buras, M. R.; Vogt, M. N.; Herasevich, V.; Dennis, J. J.; Regimbal, R. J.; Bauer, P. R.; Blair, J. E.; van Buskirk, C. M.; Winters, J. L.; Stubbs, J. R.; Paneth, N. S.; Verdun, N. C.; Marks, P.; Casadevall, A., Early safety indicators of COVID-19 convalescent plasma in 5,000 patients. The Journal of clinical investigation 2020.

10. Duan, K.; Liu, B.; Li, C.; Zhang, H.; Yu, T.; Qu, J.; Zhou, M.; Chen, L.; Meng, S.; Hu, Y.; Peng, C.; Yuan, M.; Huang, J.; Wang, Z.; Yu, J.; Gao, X.; Wang, D.; Yu, X.; Li, L.; Zhang, J.; Wu, X.; Li, B.; Xu, Y.; Chen, W.; Peng, Y.; Hu, Y.; Lin, L.; Liu, X.; Huang, S.; Zhou, Z.; Zhang, L.; Wang, Y.; Zhang, Z.; Deng, K.; Xia, Z.; Gong, Q.; Zhang, W.; Zheng, X.; Liu, Y.; Yang, H.; Zhou, D.; Yu, D.; Hou, J.; Shi, Z.; Chen, S.; Chen, Z.; Zhang, X.; Yang, X., Effectiveness of convalescent plasma therapy in severe COVID-19 patients. Proc Natl Acad Sci U S A 2020, 117, (17), 9490–9496.

11. Hurlburt, N. K.; Wan, Y.-H.; Stuart, A. B.; Feng, J.; McGuire, A. T.; Stamatatos, L.; Pancera, M., Structural basis for potent neutralization of SARS-CoV-2 and role of antibody affinity maturation. bioRxiv 2020, 2020.06.12.148692.

12. Seydoux, E.; Homad, L. J.; MacCamy, A. J.; Parks, K. R.; Hurlburt, N. K.; Jennewein, M. F.; Akins, N. R.; Stuart, A. B.; Wan, Y.-H.; Feng, J.; Nelson, R. E.; Singh, S.; Cohen, K. W.; McElrath, M. J.; Englund, J. A.; Chu, H. Y.; Pancera, M.; McGuire, A. T.; Stamatatos, L., Characterization of neutralizing antibodies from a SARS-CoV-2 infected individual. bioRxiv 2020, 2020.05.12.091298.

13. Beaudoin-Bussières, G.; Laumaea, A.; Anand, S. P.; Prévost, J.; Gasser, R.; Goyette, G.; Medjahed, H.; Perreault, J.; Tremblay, T.; Lewin, A.; Gokool, L.; Morrisseau, C.; Bégin, P.; Tremblay, C.; Martel-Laferrière, V.; Kaufmann, D. E.; Richard, J.; Bazin, R.; Finzi, A., Decline of humoral responses against SARS-CoV-2 Spike in convalescent individuals. 2020, 2020.07.09.194639.

14. Prévost, J.; Gasser, R.; Beaudoin-Bussières, G.; Richard, J.; Duerr, R.; Laumaea, A.; Anand, S. P.; Goyette, G.; Ding, S.; Medjahed, H.; Lewin, A.; Perreault, J.; Tremblay, T.; Gendron-Lepage, G.; Gauthier, N.; Carrier, M.; Marcoux, D.; Piché, A.; Lavoie, M.; Benoit, A.; Loungnarath, V.; Brochu, G.; Desforges, M.; Talbot, P. J.; Gould Maule, G. T.; Côté, M.; Therrien, C.; Serhir, B.; Bazin, R.; Roger, M.; Finzi, A., Cross-sectional evaluation of humoral responses against SARS-CoV-2 Spike. bioRxiv 2020, 2020.06.08.140244.

15. Robbiani, D. F.; Gaebler, C.; Muecksch, F.; Lorenzi, J. C. C.; Wang, Z.; Cho, A.; Agudelo, M.; Barnes, C. O.; Gazumyan, A.; Finkin, S.; Hagglof, T.; Oliveira, T. Y.; Viant, C.; Hurley, A.; Hoffmann, H. H.; Millard, K. G.; Kost, R. G.; Cipolla, M.; Gordon, K.; Bianchini, F.; Chen, S. T.; Ramos, V.; Patel, R.; Dizon, J.; Shimeliovich, I.; Mendoza, P.; Hartweger, H.; Nogueira, L.; Pack, M.; Horowitz, J.; Schmidt, F.; Weisblum, Y.; Michailidis, E.; Ashbrook, A. W.; Waltari, E.; Pak, J. E.; Huey-Tubman, K. E.; Koranda, N.; Hoffman, P. R.; West, A. P., Jr.; Rice, C. M.; Hatziioannou, T.; Bjorkman, P. J.; Bieniasz, P. D.; Caskey, M.; Nussenzweig, M. C., Convergent antibody responses to SARS-CoV-2 in convalescent individuals. Nature 2020.

16. Yuan, M.; Wu, N. C.; Zhu, X.; Lee, C. D.; So, R. T. Y.; Lv, H.; Mok, C. K. P.; Wilson, I. A., A highly conserved cryptic epitope in the receptor-binding domains of SARS-CoV-2 and SARS-CoV. Science 2020.

17. Liu, L.; Wang, P.; Nair, M. S.; Yu, J.; Rapp, M.; Wang, Q.; Luo, Y.; Chan, J. F.; Sahi, V.; Figueroa, A.; Guo, X. V.; Cerutti, G.; Bimela, J.; Gorman, J.; Zhou, T.; Chen, Z.; Yuen, K. Y.; Kwong, P. D.; Sodroski, J. G.; Yin, M. T.; Sheng, Z.; Huang, Y.; Shapiro, L.; Ho, D. D., Potent neutralizing antibodies against multiple epitopes on SARS-CoV-2 spike. Nature 2020, 584, (7821), 450–456.

18. Rogers, T. F.; Zhao, F.; Huang, D.; Beutler, N.; Burns, A.; He, W. T.; Limbo, O.; Smith, C.; Song, G.; Woehl, J.; Yang, L.; Abbott, R. K.; Callaghan, S.; Garcia, E.; Hurtado, J.; Parren, M.; Peng, L.; Ramirez, S.; Ricketts, J.; Ricciardi, M. J.; Rawlings, S. A.; Wu, N. C.; Yuan, M.; Smith, D. M.; Nemazee, D.; Teijaro, J. R.; Voss, J. E.; Wilson, I. A.; Andrabi, R.; Briney, B.; Landais, E.; Sok, D.; Jardine, J. G.; Burton, D. R., Isolation of potent SARS-CoV-2 neutralizing antibodies and protection from disease in a small animal model. Science 2020, 369, (6506), 956–963.

19. Ding, S.; Gasser, R.; Gendron-Lepage, G.; Medjahed, H.; Tolbert, W. D.; Sodroski, J.; Pazgier, M.; Finzi, A., CD4 Incorporation into HIV-1 Viral Particles Exposes Envelope Epitopes Recognized by CD4-induced Antibodies. J Virol 2019.

20. Hoffmann, M.; Kleine-Weber, H.; Schroeder, S.; Kruger, N.; Herrler, T.; Erichsen, S.; Schiergens, T. S.; Herrler, G.; Wu, N. H.; Nitsche, A.; Muller, M. A.; Drosten, C.; Pohlmann, S., SARS-CoV-2 Cell Entry Depends on ACE2 and TMPRSS2 and Is Blocked by a Clinically Proven Protease Inhibitor. Cell 2020, 181, (2), 271–280 e8.

21. Kassa, A.; Finzi, A.; Pancera, M.; Courter, J. R.; Smith, A. B., 3rd; Sodroski, J., Identification of a Human Immunodeficiency Virus (HIV-1) Envelope Glycoprotein Variant Resistant to Cold Inactivation. J Virol 2009.

22. Desormeaux, A.; Coutu, M.; Medjahed, H.; Pacheco, B.; Herschhorn, A.; Gu, C.; Xiang, S. H.; Mao, Y.; Sodroski, J.; Finzi, A., The highly conserved layer-3 component of the HIV-1 gp120 inner domain is critical for CD4-required conformational transitions. J Virol 2013, 87, (5), 2549–62.

23. Finzi, A.; Xiang, S. H.; Pacheco, B.; Wang, L.; Haight, J.; Kassa, A.; Danek, B.; Pancera, M.; Kwong, P. D.; Sodroski, J., Topological layers in the HIV-1 gp120 inner domain regulate gp41 interaction and CD4-triggered conformational transitions. Mol Cell 2010, 37, (5), 656–67.

24. Pacheco, B.; Alsahafi, N.; Debbeche, O.; Prevost, J.; Ding, S.; Chapleau, J. P.; Herschhorn, A.; Madani, N.; Princiotto, A.; Melillo, B.; Gu, C.; Zeng, X.; Mao, Y.; Smith, A. B., 3rd; Sodroski, J.; Finzi, A., Residues in the gp41 Ectodomain Regulate HIV-1 Envelope Glycoprotein Conformational Transitions Induced by gp120-Directed Inhibitors. J Virol 2017, 91, (5).

25. Long, Q. X.; Tang, X. J.; Shi, Q. L.; Li, Q.; Deng, H. J.; Yuan, J.; Hu, J. L.; Xu, W.; Zhang, Y.; Lv, F. J.; Su, K.; Zhang, F.; Gong, J.; Wu, B.; Liu, X. M.; Li, J. J.; Qiu, J. F.; Chen, J.; Huang, A. L., Clinical and immunological assessment of asymptomatic SARS-CoV-2 infections. Nat Med 2020.

26. Seow, J.; Graham, C.; Merrick, B.; Acors, S.; Steel, K. J. A.; Hemmings, O.; O’Bryne, A.; Kouphou, N.; Pickering, S.; Galao, R.; Betancor, G.; Wilson, H. D.; Signell, A. W.; Winstone, H.; Kerridge, C.; Temperton, N.; Snell, L.; Bisnauthsing, K.; Moore, A.; Green, A.; Martinez, L.; Stokes, B.; Honey, J.; Izquierdo-Barras, A.; Arbane, G.; Patel, A.; OConnell, L.; O Hara, G.; MacMahon, E.; Douthwaite, S.; Nebbia, G.; Batra, R.; Martinez-Nunez, R.; Edgeworth, J. D.; Neil, S. J. D.; Malim, M. H.; Doores, K., Longitudinal evaluation and decline of antibody responses in SARS-CoV-2 infection. 2020, 2020.07.09.20148429.

